# Pathology of Influenza A (H5N1) infection in pinnipeds reveals novel tissue tropism and vertical transmission

**DOI:** 10.1101/2025.02.07.636856

**Authors:** Carla Fiorito, Ana Colom, Antonio Fernández, Paula Alonso-Almorox, Marisa Andrada, Daniel Lombardo, Eva Sierra

**Affiliations:** Consejo Nacional de Investigaciones Científicas y Técnicas (CONICET), Argentina; Centro para el Estudio de Sistemas Marinos (CESIMAR-CONICET), Chubut, Argentina; Veterinary Histology and Pathology, Atlantic Center for Cetacean Research (CAIC), University Institute of Animal Health and Food Safety (IUSA), Veterinary School, University of Las Palmas de Gran Canaria. Trasmontaña s/n, 35413 Arucas. Canary Islands, Spain; Universidad de Buenos Aires, Facultad de Ciencias Veterinarias. Instituto de Investigación y Tecnología en Reproducción Animal (INITRA), Buenos Aires, Argentina

## Abstract

In 2023, an unprecedented outbreak of highly pathogenic avian influenza (HPAI) H5N1 resulted in the death of thousands of pinnipeds along the Argentinean coast, raising concerns about its ecological and epidemiological impact. Here, we present clinical, pathological, and molecular findings associated with HPAI H5N1 infection in pinnipeds from Chubut, Argentina. Necropsies were conducted on three South American Sea Lions (SASLs) (*Otaria flavescens*) and one Southern Elephant Seal (SES) (*Mirounga leonina*), followed by histopathological, immunohistochemical and RT-sqPCR analyses. Neurological clinical signs were observed in two SASLs, with one also exhibiting respiratory distress. Neuropathological findings included lymphoneutrophilic meningoencephalomyelitis and choroiditis, neuronal necrosis, gliosis, hemorrhages, and perivascular cuffing. Viral antigen was localized in neurons, glial cells, choroid plexus epithelial cells, ependymal cells, and the neuropil. Systemic manifestations included HPAI-related necrotizing myocarditis in the elephant seal and placental necrosis in a sea lion, with fetal tissues testing positive for HPAIV. Pulmonary lesions were minimal, limited to bronchial glands in one individual. RT-sqPCR confirmed HPAI H5 in all tested animals. Our findings highlight the neurotropism of HPAI H5N1 in pinnipeds, and expand the known systemic effects of the virus, revealing new tissue tropism and vertical transmission.

## INTRODUCTION

The highly pathogenic avian influenza virus (HPAIV) H5N1 clade 2.3.4.4b emerged in 2020 and became the predominant variant worldwide, causing intercontinental epizootics, severely impacting the poultry industry but also affecting a variety of wild or captive birds and mammals, as well as humans (1, 2). In 2021, the virus spread from Europe to North America through migratory birds (3) and reached South America by late 2022, causing high mortality events in wild birds along the Pacific coast (4, 5, 6). By the beginning of 2023, the virus spilled over into South American sea lions (SASLs) (*Otaria flavescens*), leading to a massive mortality event in Peru and Chile (7, 8).

In August 2023, the Argentinian National Service of Health and Food Quality (SENASA) reported the first cases of HPAI H5N1 in SASLs from Tierra del Fuego (6). Over the following weeks, the virus rapidly propagated along the Argentinian coast, resulting in an unprecedented mortality rate among SASLs and Southern Elephant Seals (SESs) (*Mirounga leonina*) (9, 10, 11,12). Suspected cases were confirmed by identifying clinical signs of the disease and molecular detection of the virus in biological samples from deceased animals (10,11,12). Affected SASLs showed clinical manifestations consistent with those reported in Peru and Chile (7,8), including disorientation, abnormal postures, ataxia, incoordination, total or partial paralysis of the limbs, myoclonus, nystagmus, seizures, dyspnea, abdominal breathing and profuse nasal and oral discharge. Furthermore, a high number of abortions was documented in the provinces of Rio Negro and Chubut, presumably associated with the outbreak, as some aborting females showed neurologic clinical signs (13), and one fetus tested positive by RT-PCR (11). In contrast, clinical manifestations in SESs have been scarcely documented, with reported signs including lethargy, impaired mobility, laboured breathing, nasal discharge, repetitive head or flipper movements, and tremors (12).

Influenza A virus (IAV) infection and associated disease have been documented in numerous pinniped species over the past 40 years (14), with cases predominantly linked to respiratory clinical signs and pneumonia (15,16,17,18). Recently, the HPAI H5 subtype has been associated with neurological clinical signs, acute encephalitis, and/or meningitis in pinnipeds, confirming the neurotropism of the virus (7,8,11,19,20,21,22). The virus has also been linked to acute necrotizing inflammation affecting multiple organs in 3 species of Canadian seals (22). Despite the large number of affected SASLs and SESs during the South American outbreak, research on the pathology of infection with recent HPAI H5 viruses in these species remains limited (8).

This study reports the histopathological, immunohistochemical, and molecular findings in SASLs and a SES stranded during the HPAIV H5N1 outbreak in Chubut, Argentina, providing novel insights into the viral tissue tropism and the transmission routes in pinnipeds.

## MATERIALS AND METHODS

### Sampling protocol

During the HPAIV H5N1 outbreak, Argentinean authorities implemented a strict protocol for stranded pinnipeds to prevent virus spread and protect public health. Dead animals were promptly buried *in situ* or removed from beaches, limiting full necropsies to be performed in authorized remote areas. When full necropsies were not permitted, partial sampling of the central nervous system (CNS) and lungs was conducted through the foramen magnum and thoracic windows, respectively. The samples included in this study were collected by trained veterinarians under strict biosafety protocols and with exceptional permits from Dirección de Fauna y Flora Silvestre del Chubut (Authorization N° 10/2023, DFyFSC).

### Case description and sample collection

Between August and October 2023, three SASL specimens and one SES underwent total or partial necropsy and were sampled for histopathology, immunohistochemistry, and molecular analysis (Table 1). Clinical signs were monitored in two animals that were found alive. The first was an adult SASL male identified as NEC 83, reported on August 29^th^ on a public beach near to Puerto Madryn city. The animal displayed difficulty swimming and maintaining buoyancy. After stranding, it exhibited severe dyspnea, nasal and oral discharge, and progressive neurological signs, including neck stiffness, abnormal postures, ataxia, facial twitching, body tremors, and stupor (Supplementary video S1). The animal died on August 30^th^ and was partially sampled immediately before being buried *in situ*. The second animal was an adult SASL female identified as NEC 85, reported on September 15^th^ in Punta Loma Reserve, exhibiting disorientation, abnormal postures, incoordination, and stumbling (Supplementary video S2). Over the following 3 days, the neurological signs progressed to ataxia, body tremors, and stupor, with no respiratory symptoms observed. The animal died on September 18^th^, and a complete necropsy was performed on September 21^st^ during which samples were collected. Brain samples, however, were obtained through the foramen magnum.

The remaining animals were found dead, so no clinical signs were reported. NEC 86 was an adult SASL female discovered on September 29^th^ at Cerro Avanzado public beach. A partial sampling was performed *in situ* before removing the carcass. NEC 94 was a pre-moulting male SES pup reported dead on October 11^th^ in Isla de los Pájaros Natural Reserve. A complete necropsy was performed *in situ* on October 12^th^, and the carcass was buried after sampling.

**Table 1:**
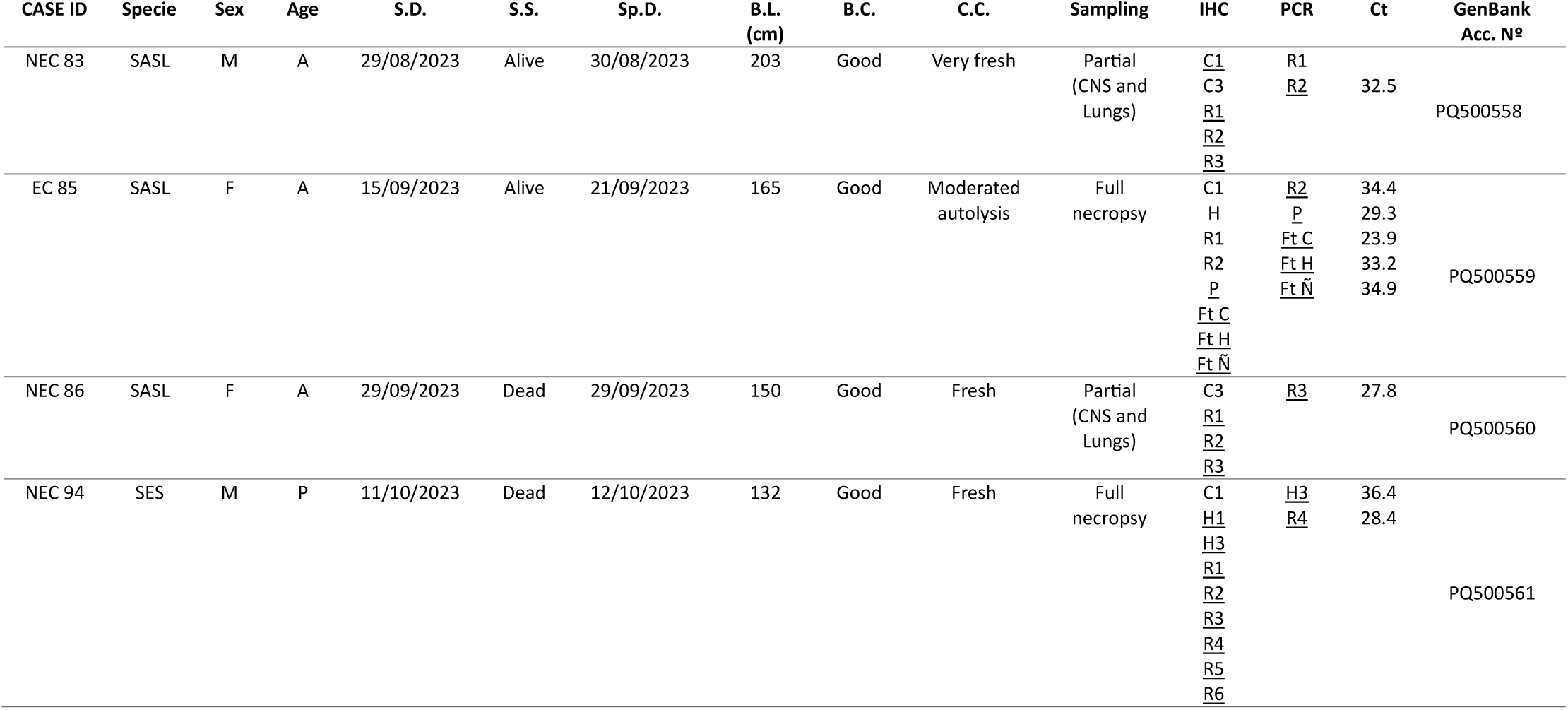
Summarized data from the animals included in the present study. Age (A = adult, P = pup); Sex (F = female, M = male); S.D. (stranding date); S.S. (stranding stage: A = alive; D = dead); S.L. (stranding location); Sp.D. (Sampling date); B.L. (Body length); B.C.: Body Condition; C.C. (carcass condition). SASL: South American Sea Lion; SES: Southern Elephant Seal; CNS: central nervous system; IHC and PCR: tested positive samples are underlined; Ct: Cycle threshold; and GenBank Acc. N°: GenBank accession number. Samples code: NEC 83: C1-C3: lungs; R1-R2: Brainstem; R3: Cerebellum and spinal cord. NEC 85: C1: lung; H: heart; R1-R2: Brainstem and medulla oblongata. NEC 86 C3: Lungs; R1: Hippocampus and choroid plexus; R2: spinal cord; R3: Brainstem, choroid plexus, and cerebellum; R6: deep white matter of the cerebellum. P: placenta. Ft: fetus. Ft C: fetal lung; Ft H: fetal heart; Ft Ñ: fetal kidney and thymus. NEC 94: C1: Lungs; H1-H3: Heart; R1-R3: choroid plexus and cerebellum; R4-R6: cerebral cortex.

Gross lesions and additional data including body length, age category, and body condition were documented. Age category was estimated based on external features including body length, fur color, and tooth examination. Body condition was assessed through inspection of subcutaneous fat during sampling. Carcasses decomposition status was classified as ‘very fresh’, ‘fresh’, ‘moderate autolysis’, ‘advanced autolysis’, or ‘very advanced autolysis’, following the classification system by Geraci (23).

### Histopathology and Immunohistochemistry

During necropsies, formalin-fixed samples of CNS and lungs were systematically collected from each animal. Additionally, samples of heart, kidneys, liver, lymph nodes, and spleen were collected from NEC 85 and NEC 94. The placenta, umbilical cord, and fetal organs, including the lungs, heart, thymus, liver and kidneys, were also sampled from specimen NEC 85. All tissue samples were processed following standard histological protocols, embedded in paraffin, sectioned at 5 μm thickness, and stained with hematoxylin and eosin (HE) for microscopic examination (24). Immunohistochemistry (IHC) was subsequently performed on 3 μm-thick sections of Formalin-Fixed Paraffin-Embedded (FFPE) tissues, specifically focusing on regions with lesions suggestive of HPAI infection (Table 1). A monoclonal antibody (EBS-I-238; Biologicals Limited, https://biologicals-ltd.com) targeting the nucleoprotein of influenza virus type A was used as previously described (8). Briefly, sections were deparaffinized, rehydrated, and subjected to antigen retrieval with ready-to-use (RTU) proteinase K for 6 minutes at room temperature (RT). Endogenous peroxidase activity was blocked using the EnVision FLEX Mini Kit (High pH, Dako) for 10 min at RT. The primary antibody was diluted to 1.5 μg/ml in RTU reagent (EnVision FLEX Antibody Diluent) and incubated for 30 min at RT, followed by incubation with an indirect peroxidase polymeric detection kit (EnVision FLEX Mini Kit, High pH, Dako) for 30 minutes at RT. The reaction was developed with Magenta solution (HRP Magenta, Dako). Slides were counterstained with Mayer’s hematoxylin, coverslipped, and examined by light microscopy (Olympus BX51, Tokyo, Japan). Imaging was performed using Camera software for DP21 (Version 02.01.01.93) (Olympus DP21, Tokyo, Japan). A positive control (brain tissue known to react positively with this specific monoclonal antibody and confirmed molecularly positive for H5N1) and negative control (the same brain section with the primary antibody omitted) were included.

### Molecular analysis

Molecular techniques were conducted on selected CNS FFPE samples, as brain tissue is considered the optimal diagnostic specimen for confirming HPAI infection (25, 26) (Table 1). The heart from NEC 94 and the placenta and fetal organs from NEC 85 were also analyzed to confirm the involvement of the HPAIV H5 in the observed lesions, as these organs had not been previously reported to be infected in pinnipeds. Four 10 μm thick sections of each FFPE sample were used for RNA extraction with the Qiagen RNeasy FFPE Kit (Qiagen, Inc., Valencia, CA, USA). Xylene deparaffinization was performed twice at 50 °C for 3 min each, followed by two washes with ethanol after removal of residual xylene and overnight digestion at 37 °C.

The molecular detection of avian influenza Hemagglutinin (HA) subtype H5 was performed using an RT-sqPCR as previously described (27). PCR product purification was performed with the Real Clean Spin kit (REAL®, Durviz, S.L., Valencia, Spain) for bidirectional sequencing via the Sanger method. The obtained sequences were aligned using the ClustalW algorithm in MEGA11 software (Pennsylvania, PA, United States) (28, 29) to generate a consensus sequence. A BLAST search was then conducted to confirm the identity of the PCR amplicon by comparing it with similar sequences in GenBank. In addition, data analysis was carried out using FluSurver (Bioinformatics Institute, A*STAR, https://flusurver.bii.a-star.edu.sg/).

## RESULTS

The data for each specimen, including case ID, species, sex, age class, stranding information, sampling date, body length, carcass condition, and sampling details, are compiled in Table 1. The table also summarizes the histopathological findings and details of samples subjected to immunohistochemistry and RT-sqPCR analyses, providing a comprehensive overview of the diagnostic and molecular investigations carried out for each case.

### Gross pathology

Common findings across all individuals included a good nutritional condition, with no evidence of muscle or fat depletion, and generalized congestion, most notably in the lungs, which were markedly atelectatic and released abundant blood and reddish foam upon incision. Three specimens (NEC 83, NEC 86, and NEC 94) presented abundant reddish, high-viscosity mucus in the larynx and a mild accumulation of serosanguinous fluid in the thoracic cavity. Furthermore, NEC 83 showed multiple epidermal lacerations, likely related to the physical trauma from active stranding. Specimens NEC 85 and NEC 94 exhibited marked hemopericardium and an empty stomach. Additionally, the female NEC 85 showed generalized lymphadenomegaly and was confirmed to be pregnant, carrying a 34 cm hairless fetus. The placenta displayed multifocal orange discoloration, while the amniotic fluid remained clear with a slight reddish tint. The fetus was apparently normal, with no visible macroscopic changes. Concurrently, the CNS of NEC 83 and NEC 94 showed marked meningeal congestion and multifocal hemorrhages in both the gray and white matter of the cerebrum and cerebellum.

### Histopathology and Immunohistochemistry

All animals presented histological lesions in the CNS, that varied in severity and distribution. Samples from NEC 83, including the brainstem, cerebellum, and spinal cord, revealed moderate to severe meningitis and inflammation in both the gray and white matter, characterized by a mix of lymphocytes, plasma cells, macrophages, and neutrophils. Specifically, inflammation was observed in the meninges of all examined CNS samples, the midbrain and the pons, the inner medulla of the cerebellum, and the anterior and posterior gray commissures surrounding the central canal in the spinal cord. Additional neuropathological findings included multifocal neuronophagia, gliosis, lymphoplasmacytic perivascular cuffing, vasculitis, and hemorrhages. AIV immunopositivity was observed in scattered neurons and glial cells, predominantly associated with the inflammatory response, as well as within the neuropil of the gray matter (Figure 3). NEC 85 presented minimal brainstem lesions, characterized predominantly by lymphoneutrophilic meningitis, with no detection of AIV immunostaining in any of the analyzed samples. In NEC 86, the primary brain lesions included mild lymphoneutrophilic meningitis and choroiditis, spongiosis, gliosis, and vascular changes such as congestion and microhemorrhages in all the analyzed samples. Immunohistochemistry revealed abundant influenza virus antigen mainly in epithelial cells of the choroid plexus, ependymal cells of the ventricles, and the central canal of the spinal cord, as well as in a few neurons and glial cells (Figure 4). Additionally, multifocal immunopositivity was detected in certain inflammatory cells, primarily macrophages, within the meninges. The specimen NEC 94 showed the most severe and extensive lesions predominantly characterized by mild to moderate lymphoneutrophilic meningitis and severe multifocal areas of encephalitis with severe neuronal and glial cell degeneration and necrosis, neuronophagia, glial cell aggregates and perivascular cuffing along the gray matter of the cerebrum and cerebellum. Degenerated and necrotic neurons, along with associated glial cells, displayed irregular morphology, dark-stained nuclei (eosinophilic or basophilic), and prominent pericellular halos. Nuclear features also included a marked reduction in euchromatin and diminished or absent nucleoli. Influenza virus antigen was predominantly detected within necrotic and inflammatory lesions, specifically within the cytoplasm and nuclei of degenerated neurons and glial cells, with intense immunostaining in hyperchromatic nuclei. Additionally, the antigen was localized in the choroid plexus epithelial cells and ependymal cells.

**Figure 1.**
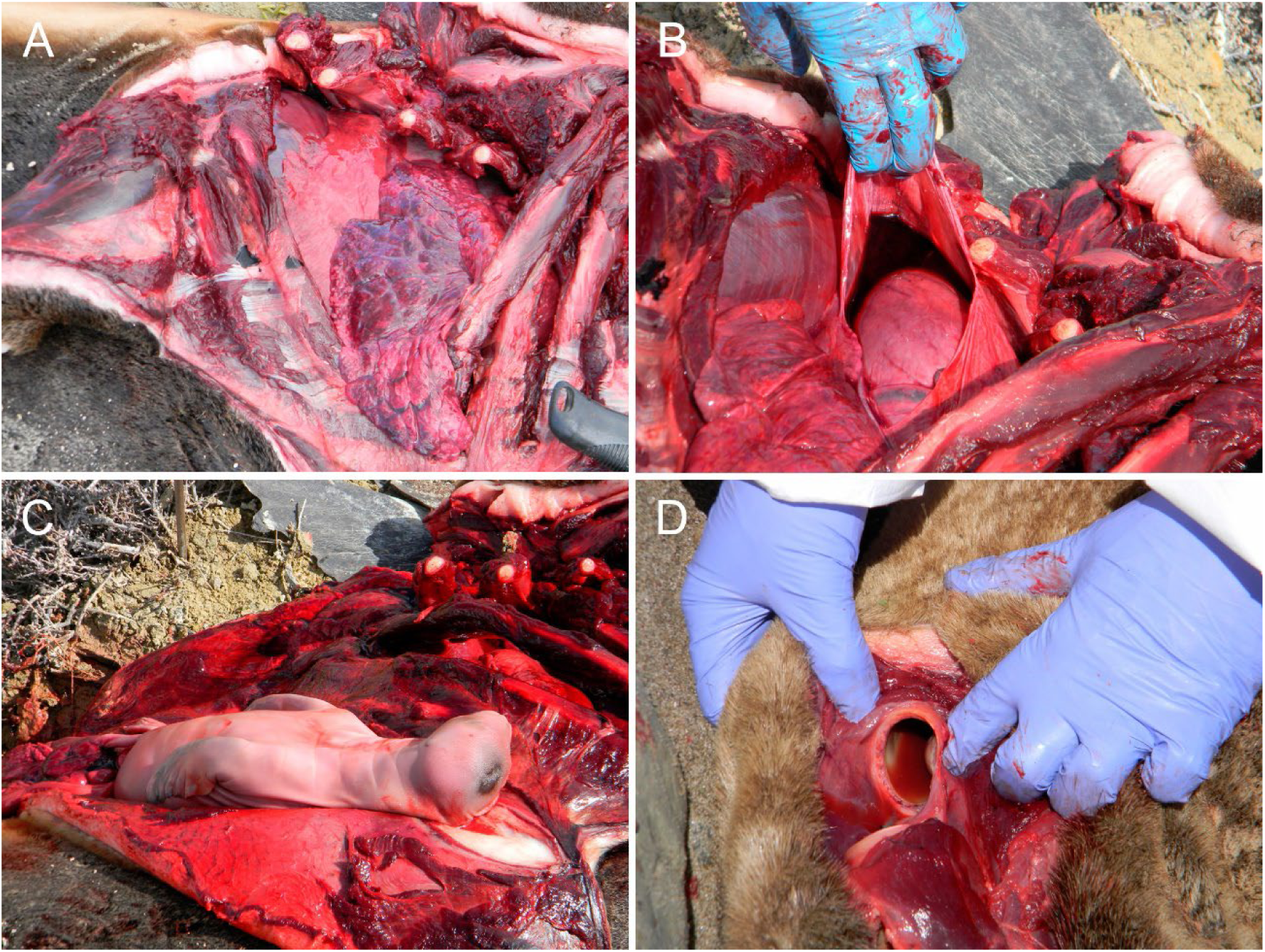
Postmortem findings in South American Sea Lions infected with highly pathogenic avian influenza A(H5N1) virus. The presence of abundant subcutaneous fat indicates good nutritional condition. A) Lungs displaying generalized congestion and severe atelectasis (NEC 85). B) Marked hemopericardium (NEC 85). C) Hairless fetus (NEC 85). D) Presence of reddish fluid in the tracheal lumen (NEC 86).

**Figure 2.**
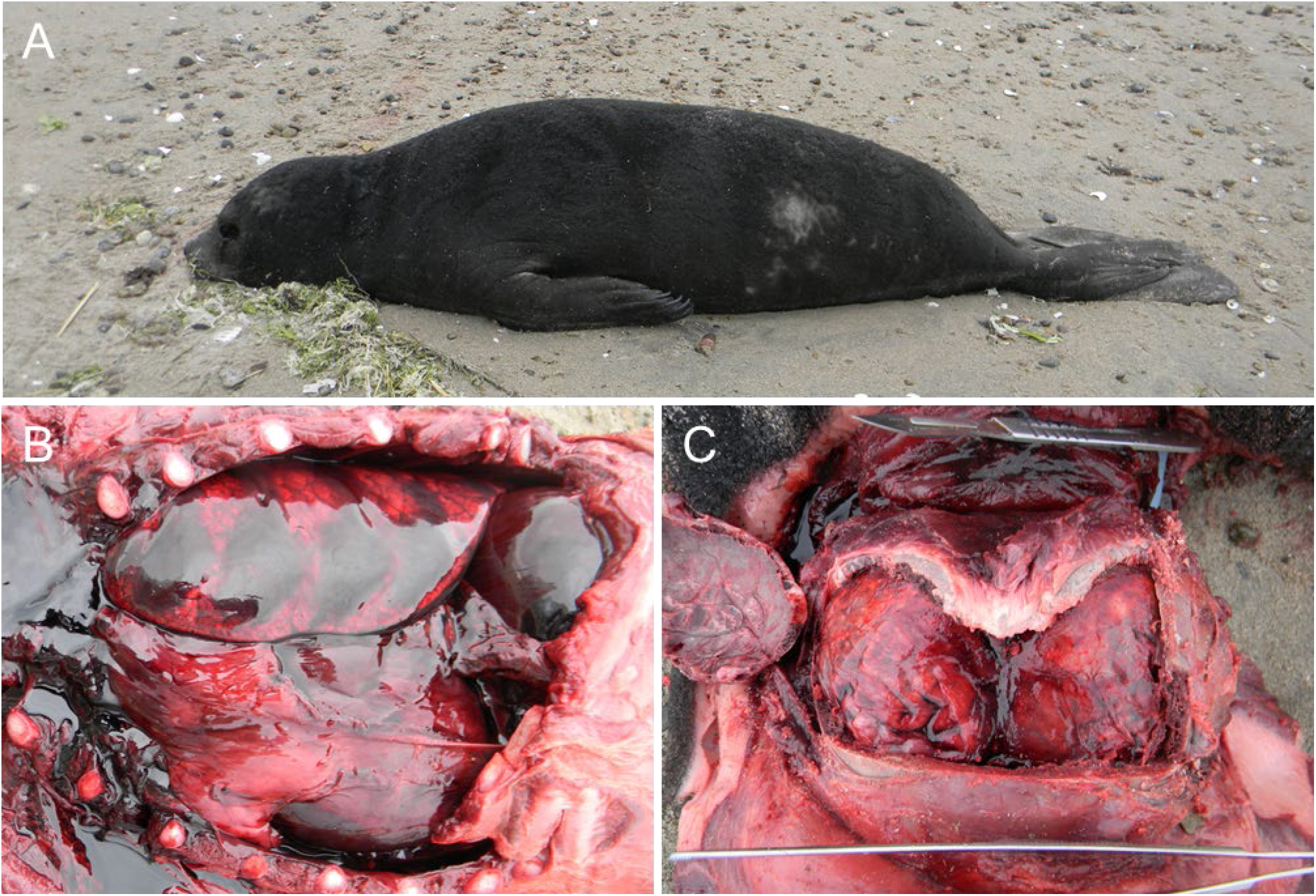
Postmortem findings in a Southern Elephant Seal pup (NEC 94), infected with highly pathogenic avian influenza A(H5N1) virus. A) The dark fur indicates a pre-moulting pup. B) Diffuse marked edematous lungs with multifocal congestive-hemorrhagic (hypostatic) areas and reddish fluid in the thoracic cavity (hemothorax). C) Marked meningeal congestion and edematous brain.

**Figure 3.**
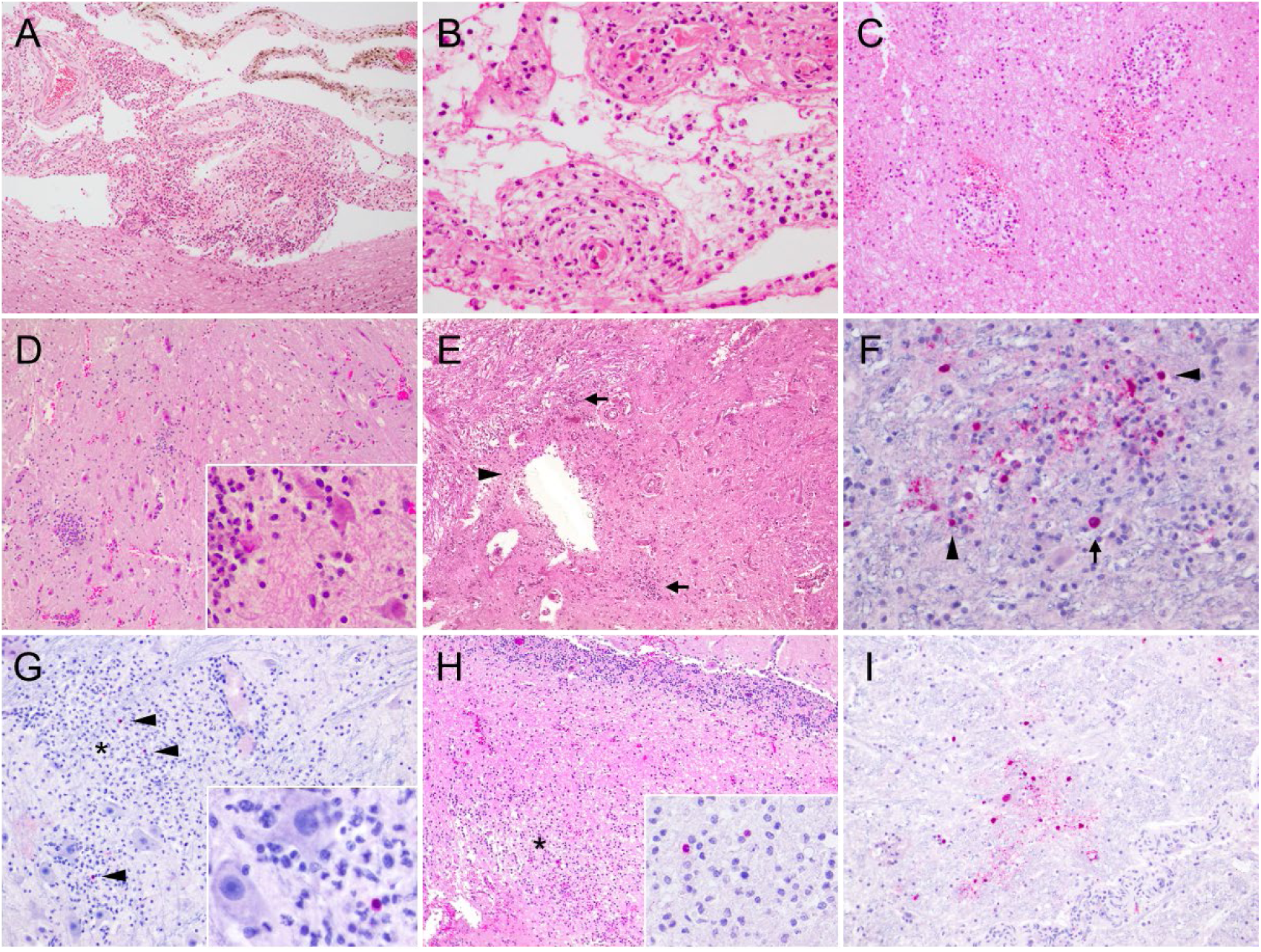
Histopathological and immunohistochemical findings in the central nervous system of South American Sea Lion NEC 83, infected with HPAI H5N1. A) Sample R1 (brainstem): severe meningitis with marked congestion and hemorrhages. H&E, 10X. B) Sample R1 (brainstem): meningitis and necrotizing vasculitis affecting small vessels (arrows). Inflammatory infiltrates are predominantly composed of mixed lymphoplasmacytic cells and neutrophils. H&E, 40X. C) Sample R3 (cerebellum): multifocal hemorrhages in the deep cerebellar white matter with mononuclear perivascular infiltrates. H&E, 20X. D) Sample R2 (brainstem): multifocal lymphoneutrophilic encephalitis, microhemorrhages, neuronal necrosis, and satellitosis. H&E, 10X. Inset: higher magnification showing neuronal necrosis, satellitosis, and neuronophagia. H&E, 40X. E) Sample R3 (spinal cord): multifocal lymphoneutrophilic myelitis (arrows), hemorrhages, and congestion. Loss of the ependymal epithelial layer is noted, with mononuclear inflammatory cells surrounding the central canal (arrowhead). H&E, 10X. F) Sample R2 (brainstem): a focus of encephalitis showing AIV-immunopositivity with signals in neuronal nuclei (arrows) and glial cells (arrowheads). IHC against Influenza A nucleoprotein, 40X. G) Sample R1 (brainstem): focal extensive focus of encephalitis (asterisk) showing neuronophagia and perivascular cuffing, with occasional intralesional AIV-immunopositive glial cells. IHC against Influenza A nucleoprotein, 20X. Inset: higher magnification of an AIV-immunopositive glial cell. IHC against Influenza A nucleoprotein, 60X. H) Sample R3 (cerebellum): focal extensive inflammatory infiltration in the cerebellar inner medulla (asterisk) consisting of lymphocytes, plasma cells, macrophages, and neutrophils. H&E 10X. Inset: AIV-immunopositive glial cell within the same lesion. IHC against Influenza A nucleoprotein, 60X. I) Sample R2 (brainstem): AIV-immunopositive neurons and glial cells without associated inflammatory reaction. IHC against Influenza A nucleoprotein, 20X.

**Figure 4.**
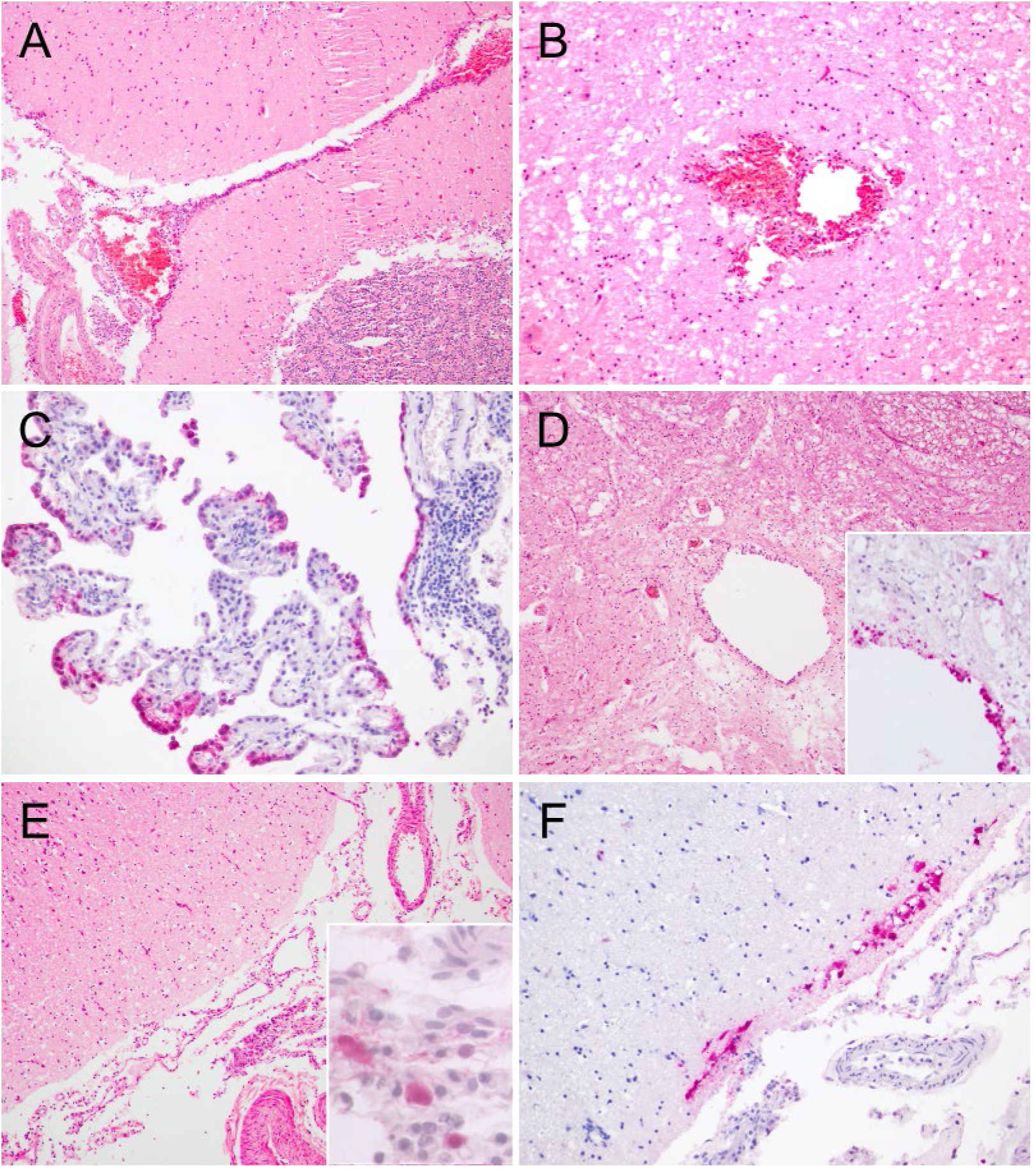
Histopathological and immunohistochemical findings in the central nervous system of South American Sea Lion NEC 86, infected with HPAI H5N1. A) Sample R3: mild lymphoneutrophilic meningitis in the cerebellum. H&E, 10X. B) Sample R6: spongiosis and micro hemorrhages in the deep cerebellar white matter. H&E, 20X. C) Sample R1: non-suppurative choroiditis (asterisk) with AIV-immunopositive choroid plexus epithelial cells (arrows). IHC against Influenza A nucleoprotein, 20X. D) Sample R2: mild diffuse myelitis characterized by mononuclear cell infiltration around the central canal of the spinal cord. H&H, 10X. Inset: AIV-immunopositive staining observed in ependymal cells (arrow), glial cells, and neural tissue surrounding the central canal. IHC against Influenza A nucleoprotein, 20X. E) Sample R1: mild lymphoneutrophilic meningitis in the entorhinal cortex. H&E, 10X. Inset: AIV-immunopositive staining observed in mononuclear inflammatory cells (macrophages), within the meninges. IHC against Influenza A nucleoprotein, 60X. F) Sample R1: AIV-immunopositive glial cells and neuropil in the gray matter of the entorhinal cortex. IHC against Influenza A nucleoprotein, 20X.

The heart of NEC 94 exhibited mild to moderate multifocal myocardial necrosis, lymphoplasmacytic inflammation, and calcification. Intralesional AIV antigens were detected in both the nucleus and cytoplasm of cardiomyocytes, confirming that these lesions were associated with HPAIV infection (Figure 6). NEC 85 showed moderate to severe lymphoplasmacytic myocarditis along with acute degenerative changes, including contraction band necrosis, hypereosinophilia, and cytoplasmic vacuolization. No AIV-immunostaining was detected in the cardiac tissue of this animal, so these changes could be likely related to stress rather than the direct effects of the virus.

**Figure 5.**
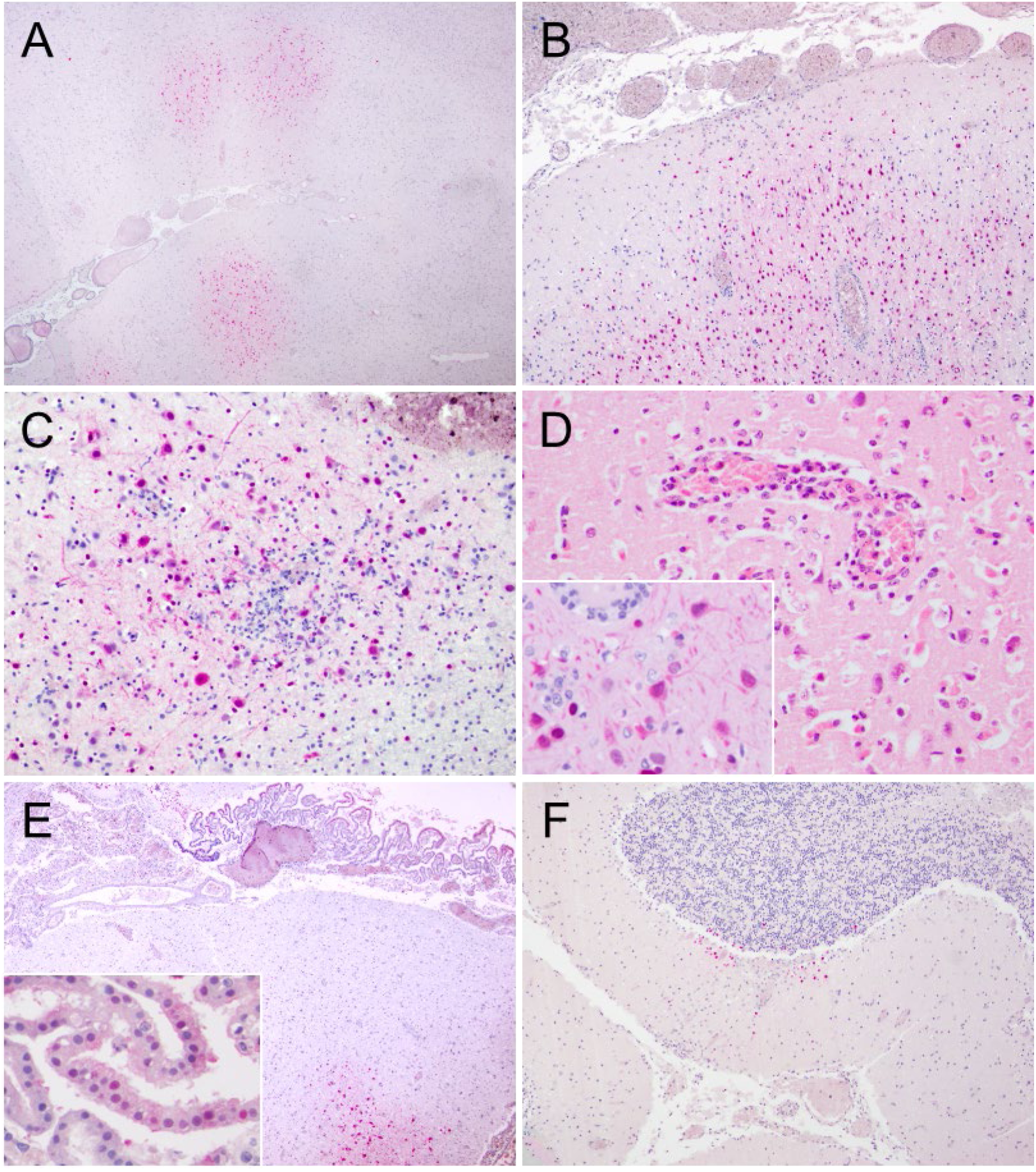
Histopathological and immunohistochemical findings in the central nervous system of Southern Elephant Seal NEC94, infected with HPAI H5N1. A) Sample R6: multifocal areas of AIV-immunopositivity observed in the gray matter of the cerebral cortex. IHC against Influenza A nucleoprotein, 0.73X. B) Sample R4: Numerous AIV-immunopositive neurons and glial cells within multifocal areas of the cortex, accompanied by perivascular cuffing and mild lymphoneutrophilic meningitis. IHC against Influenza A nucleoprotein, 10X. C) Sample R4: lymphoneutrophilic encephalitis with intralesional AIV-immunopositive neurons and glial cells. IHC against Influenza A nucleoprotein, 20X. D) Sample R6: perivascular lymphohistiocytic infiltration, neuronal necrosis, and inflammatory cell infiltration in the gray matter. H&E, 40X. Inset: abundant AIV nucleoprotein viral antigen detected in the nuclei and cytoplasm of neurons and glial cells. IHC against Influenza A nucleoprotein, 40X. E) Sample R2: lymphoneutrophilic choroiditis (asterisk) with extensive AIV-immunopositivity in the choroid plexus epithelial cells, as well as in neurons and glial cells associated with focal encephalitis foci. IHC against Influenza A nucleoprotein, 4X. Inset: higher magnification of AIV-immunopositive choroid plexus epithelial cells. IHC against Influenza A nucleoprotein, 40X. F) Sample R1: mild lymphoneutrophilic meningitis in the cerebellum with scattered AIV-immunopositive glial cells in the glomerular zone, molecular layer, and around Purkinje cells. IHC against Influenza A nucleoprotein, 10X.

**Figure 6.**
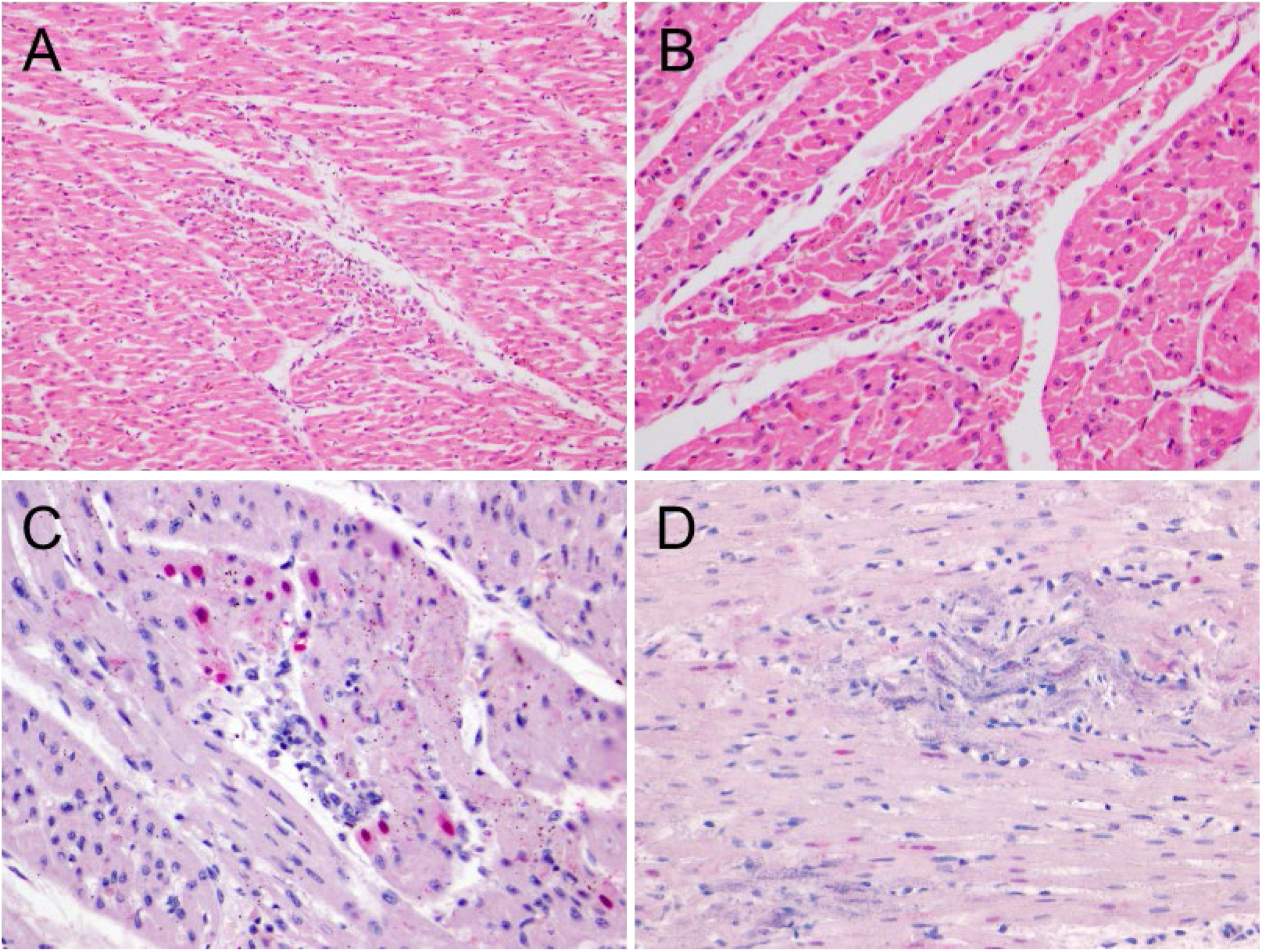
Histopathological and immunohistochemical findings in the heart of Southern Elephant Seal NEC 94, infected with HPAI H5N1. A) Focal myocardial necrosis with inflammatory infiltrate. H&E,20X. B) Perivascular inflammatory infiltrates with mild cardiomyocyte degeneration. H&E, 40X. C) Mild granulomatous myocarditis with myocardial necrosis and AIV-immunopositive nuclei in cardiomyocytes. IHC against Influenza A nucleoprotein, 40X. D) Multifocal mineral deposition (consistent with calcium) within the myocardium, associated with positive viral protein staining in the nuclei of a few cardiomyocytes. IHC against Influenza A nucleoprotein, 40X.

The placental labyrinth from NEC 85 showed multifocal areas of chorionic villi necrosis affecting the trophoblast cells and maternal vessels with positive intralesional AIV immunostaining. AIV antigens was also detected in the nucleus of apparently normal trophoblast cells, mononuclear cells with morphological features of macrophages within the core of chorionic villus, fusiform cells and macrophages in the maternal connective tissue (Figure 7). Regarding fetal organs, the lungs showed intense AIV-positive staining in epithelial cells lining embryonic airways, interstitial and luminal mononuclear cells and some luminal epithelioid cells. The thymus presented intense AIV immunopositivity in macrophages and stromal cells within the medulla, and a few lymphocytes and endothelial cells. AIV antigen was also detected in the nucleus and cytoplasm of cardiomyocytes and scattered epithelial cells in the kidney.

**Figure 7.**
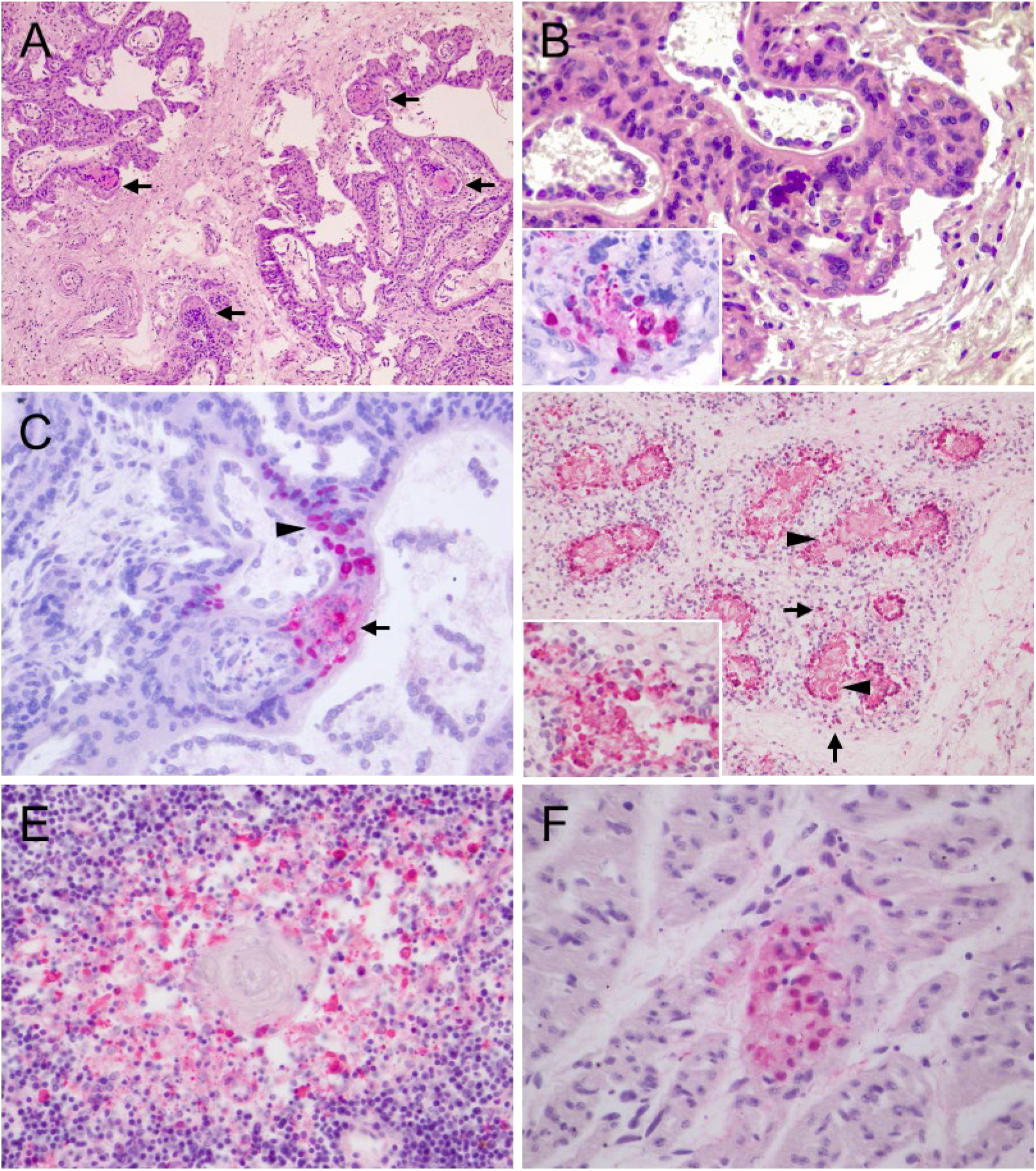
Histopathological and immunohistochemical findings in the placenta and fetus of South American Sea Lion NEC85, infected with HPAI H5N1. A) Multifocal areas of necrosis in the chorionic villi. H&E, 10X. B) Necrotic fetal chorionic villi with histiocytic inflammation. H&E, 20X. Inset: AIV-immunopositivity in the nuclei and cytoplasm of large mononuclear cells with morphological features of macrophages in the core of a necrotic chorionic villus. IHC against Influenza A nucleoprotein, 40X. C) Intranuclear AIV-immunostaining was observed in trophoblastic cells within necrotic lesions (arrow) and normal trophoblastic cells (arrowheads). IHC against Influenza A nucleoprotein, 40X. D) Fetal lung with intense positivity in epithelial cells lining the lumen of immature airways, interstitial macrophages (arrows), and epithelioid cells (arrowhead), 20X. Inset: abundant AIV nucleoprotein viral antigen detected in luminal mononuclear cells, 40X. IHC against Influenza A nucleoprotein. E) Fetal thymus showing abundant immunopositivity in medullary macrophages and thymic epithelial cells. IHC against Influenza A nucleoprotein, 40X.F) AIV-immunopositive in fetal cardiomyocytes. IHC against Influenza A nucleoprotein, 60X

Lung samples from the 4 animals showed mostly vascular changes including congestion, oedema, alveolar and bronchiolar hemorrhages, intravascular coagulation, and leukocytosis. The only HPAI-related lesions identified in the lungs were bronchial gland epithelial necrosis, confirmed immunohistochemically in NEC 83 (Figure 8). NEC 85 presented mild multifocal fibrinous necrotic bronchopneumonia, while NEC 94 showed moderate to severe granulomatous bronchopneumonia associated with intralesional bacteria. No AIV immunostaining was detected in lung samples from NEC 85, NEC 86, and NEC 94.

**Figure 8.**
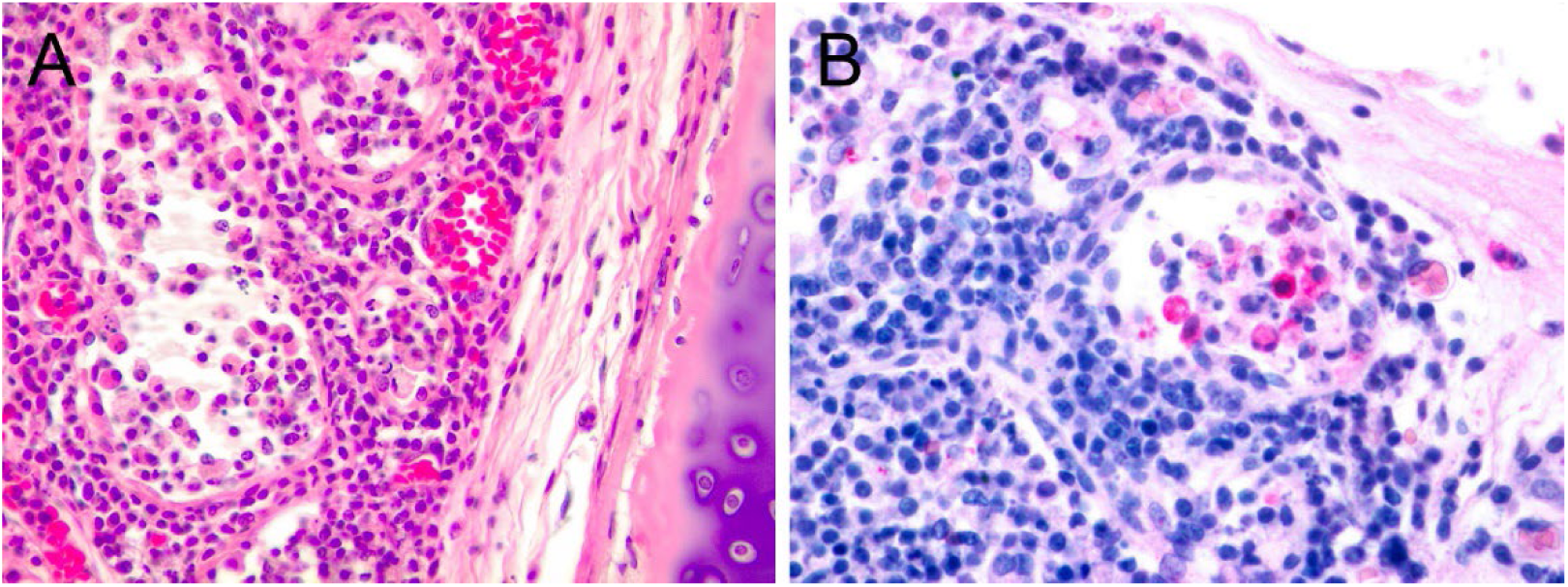
Histopathological and immunohistochemical findings in the lungs of South American Sea Lion NEC83, infected with HPAI H5N1. A) bronchial gland with necrosis of epithelial cells and associated granulomatous inflammation. H&E, 40x. B) intralesional positive viral protein staining in mononuclear and epithelial cells. IHC against Influenza A nucleoprotein, 60X.

No significant histological findings or AIV expression were detected in the remaining examined tissue samples.

### Molecular results

Ten of the 11 FFPE samples tested by RT-sqPCR were confirmed positive for HPAI H5, including NEC 83-R2, NEC 85-R2, NEC 86-R3, NEC 94-R4, NEC 94-heart, NEC 85-placenta and fetal samples of lung, heart, kidney and thymus. Amplicons of approximately 150 base pairs were successfully obtained and purified for sequencing from the remaining samples. The cycle threshold (Ct) values ranged from 23.9 to 36.4, with a mean of 31.2 and a standard deviation (SD) of 3.8 (Table 1). Two distinct sequences, each 152 base pairs in length, were identified: Sequence (SQ) 1, obtained from samples NEC 85-R2, NEC 85-placenta (PQ500559), NEC 85-fetus, NEC 86-R3 (PQ500560), NEC 94-R4 and NEC 94 H3 (PQ500561), and SQ2, from NEC 83-R2 (PQ500558). Both sequences were confirmed as HPAI H5N1 clade 2.3.4.4b when the HA segment sequences were compared with various H5N1 clades. BLAST analysis revealed that the HA sequences of SQ1 and SQ2 share 99.00% similarity. Furthermore, they exhibited 96.5% to 95.39% sequence identity and 100% query coverage (QC) with partial HA gene segments from various avian and mammalian species sampled in North and South America between 2022 and 2024. These included sequences from two SESs (PP488329, PQ002114) and a SASL (OR987092) collected in Argentina. This HA gene encodes the hemagglutinin protein, which is essential for the virus’s ability to infect host cells and plays a pivotal role in its virulence and transmissibility.

## DISCUSSION

The 2022-2023 outbreak of H5N1 HPAI clade 2.3.4.4b in South America led to an unprecedented large-scale mortality event among pinnipeds, drastically affecting the SASL populations throughout their entire geographical range (6, 7, 8). Additionally, the virus severely impacted some breeding colonies of SESs at Peninsula Valdés, Argentina, yielding mortality of 96% of the pups born during the 2023 breeding season (9, 12). This catastrophic mortality event highlights the severe ecological consequences of emerging pathogens like HPAI viruses on marine mammal populations and underscores the urgent need for long-term health surveillance actions.

The rapid spread and the high frequency of HPAIV H5N1 clade 2.3.4.4b spillover into domestic and wild mammals, alerts about the virus’s adaptability to new hosts, increasing concern regarding public health and the risk of a potential pandemic event. Molecular techniques, followed by genome sequencing, have been extensively used during the South America outbreak to characterize virus strains and detect mutations related to mammal-to-mammal transmission (7, 8, 11, 12, 31). However, knowledge about the pathogenicity and tissue tropism of the virus in pinnipeds remains scarce, with only a limited number of studies addressing this critical issue (8). Our study describes the pathology associated with HPAIV H5N1 clade 2.3.4.4b infection in three adult SASLs and one SES pup from Chubut, Argentina, revealing novel insights into viral tissue tropisms and advancing our understanding of HPAIV pathology in pinniped species. This study offers a critical addition to the current understanding of the impact of clade 2.3.4.4b AIV on non-avian wildlife, underscoring the virus’s capacity to infect diverse host tissues and the potential implication for pinniped health and conservation.

Gross examination revealed only minor, non-specific changes across all 4 analyzed animals, likely reflecting the disease’s rapid progression. All necropsied pinnipeds were found to be in good nutritional condition, indicating an acute or super-acute onset of death. This finding aligns with previous reports in pinnipeds and other mammalian species, where similar acute clinical presentations have been documented in association with HPAIV infections (8, 21, 22, 30, 32). Histological examination, however, revealed widespread lesions consistent with HPAI H5 infection in the CNS, heart, lungs, and placenta, which were confirmed by immunohistochemistry and RT-sqPCR, providing strong evidence of a causal association between HPAIV infection, clinical signs, tissue damage, and mortality in these marine mammals. We provide the first evidence of HPAIV H5N1 infection in the placenta and fetal tissues from a pregnant SASL, supporting transplacental infection and vertical transmission.

The neuroinvasive potential and the ability to replicate within the CNS causing severe neurological disease is a distinctive characteristic of HPAI H5 viruses that has been reported previously in birds and mammals, including pinnipeds and humans (30). Our study confirms the highly neurotropic nature of HPAI H5N1 in SASL and SES and its ability to induce severe lesions in multiple regions of the CNS, including the cortex, hippocampus, midbrain, choroid plexus, brainstem, cerebellum, and spinal cord. The most common lesions observed in the CNS included moderate to severe lymphoneutrophilic meningoencephalitis, neuronal necrosis, glial cell aggregates, and perivascular cuffing. These agreed with previous descriptions in pinnipeds (8, 21, 22) and provide a basis for understanding the neurological signs frequently observed in these marine mammals. A novelty finding for pinnipeds includes multifocal myelitis and choroiditis with AIV antigen detection in the epithelial cells of the choroid plexus and the ependymal cells lining the central canal of the spinal cord, suggesting that the virus spread via the cerebrospinal fluid, as has been previously suggested in mammals and birds (30, 34, 36). Experimental intranasal inoculation of Influenza A H5N1 strains caused similar CNS lesions in ferrets and mice, and demostrated that the virus initially infected cells within the nasal mucosa and reached the olfactory bulb via cranial nerves with subsequent dissemination through the cerebrospinal fluid (33, 34, 35). In ferrets, the nasal route of entry explains the neuroinvasive ability of the H5N1 virus as the weak barrier between the olfactory bulb and cerebrospinal fluid allows access to the tissues surrounding the subarachnoid space and ventricular system (33, 34). On the other hand, there is little evidence supporting neuroinvasion via hematogenous route even when HPAI H5Nx viruses can spread to the circulation (viremia) in both humans and experimentally inoculated animals (30). The transmission pathway of HPAIV H5N1 in pinnipeds has been widely debated, and molecular analyses indicate that the virus circulating in South American pinnipeds harbors mutations associated with increased virulence, mammalian host adaptation, and mammal-to-mammal transmission (12, 31). The rapid spread of the virus among colonies, and the hyperacute neurological disease strongly suggest direct contact and respiratory/nasal pathways as the most likely route of infection in these marine mammals (12, 31). Our results provide morphological cues supporting the nasal route of infection for the disease in pinnipeds. However, the precise infection route in pinnipeds remains undetermined.

When comparing corresponding regions of the CNS (cortex, brainstem, and cerebellum), lesions in the SES pup were notably more severe than those observed in the SASLs. Previous research in birds has demonstrated that different avian species exhibit varying susceptibility to HPAI H5 viruses (36), which may partially explain interspecies differences in CNS lesion severity. This variability is likely influenced by host-specific factors, such as variations in immune responses, viral tropism, and receptor distribution within the CNS. However, due to the sampling method in SASLs, which involved accessing tissue via the foramen magnum, detailed information on the distribution of lesions in these animals was limited.

The current H5N1 clade 2.3.4.4b outbreak is highlighted by the high neurotropism and neurological disease, with limited viral-related lesions or viral detection in the respiratory system of several species, including marine mammals (8, 30, 37, 38). In our study, lung lesions associated with AIV were immunohistochemically confirmed only in NEC 83, which also exhibited respiratory clinical signs. The lesions observed in NEC 83 were identical to those previously described in pinnipeds, predominantly affecting the bronchial glands (8; 22). No lung tissue samples from the remaining animals tested positive for AIV by IHC. The detection of AIV antingens only in NEC 83 confirms the lower tropism of the virus for the respiratory system. This pattern aligns with observations in other pinniped species, where neurological lesions are generally more pronounced than respiratory ones (8, 16). However, in the absence of PCR results for the lung samples, a definitive conclusion cannot be drawn, as some CNS samples that were negative by IHC tested positive by PCR. This discrepancy underscores the need for further research regarding HPAIV H5N1 impact on the respiratory system in pinnipeds.

The cardiac lesions identified in the SES (NEC 94), confirmed that the HPAI H5N1 can cause necrotizing myocarditis, providing novel information about tissue tropism of the virus in pinnipeds. Influenza-associated myocarditis has been documented in wild and domestic avian species infected with HPAI clade 2.3.4.4b (39, 40, 41, 42). In terrestrial mammals, multifocal myocardial necrosis has been reported in natural H5N1 infections (32, 39, 43) and in experimentally infected cats (44). Additionally, multifocal myocarditis was noted in a seal during the 2022 HPAI H5N1 outbreak in Canada, although confirmation by molecular or IHC methods was not performed (22). In humans, acute myocarditis is a well-recognized complication of influenza A virus subtypes H5N1 and H1N1, often resulting in severe cardiac dysfunction or death (45). Although the limited sample size precludes determining whether myocarditis is a common HPAI complication in pinnipeds, the severity and extent of cardiac lesions observed in NEC 94 suggest that myocarditis could contribute to mortality in affected individuals. Additionally, the presence of viral antigens in the cardiomyocytes of the fetus confirms the myocardial tissue tropism of the virus in SASLs, despite no associated lesions being observed in this specimen. Our findings provide new insights into the pathogenesis of the virus in pinnipeds. Therefore, HPAI H5N1 should be considered in the differential diagnosis of viral myocarditis in these species.

A significant finding of our study was the identification of placental lesions associated with HPAIV clade 2.3.4.4b in the pregnant SASL, demonstrating active viral presence and replication within placental tissues. Furthermore, fetal organs tested positive for AIV nucleoprotein and PCR, providing compelling evidence that supports the transplacental route as a mode of HPAIV H5N1 transmission. Vertical transplacental transmission of HPAIV has been previously documented in humans (46, 47) and in the BALB/c mouse model (48). Additionally, mother-to-calf transmission via milk has been recently proposed in dairy cattle (49, 50). To our knowledge, this represents the first reported case of placental and fetal HPAIV H5N1 infection in a wild mammal, offering novel insights into the virus’s pathogenesis and transmission dynamics. In addition to neurologic signs commonly observed in pinnipeds, an atypical abortion rate was reported during the HPAIV outbreak in several SALS colonies along the coast of Chubut and Río Negro, Patagonia (13). To date, only two fetuses have tested positive for the AIV by PCR across South America (7, 11), but no histopathological or IHC analyses were conducted on these specimens to confirm viral infection or cytopathic effect. Our data support that the H5N1 viruses could vertically transmit to the fetus in pinnipeds giving a possible explanation for the high number of SASL abortions recorded during the Argentina outbreak (13). Alternatively, AIV-related abortions in humans have been linked to increased production of proinflammatory cytokines by the infected placenta, rather than being directly attributable to the cytotoxic effects of the virus itself (51). This immunological response may also play a role in the abortion event observed in the SASL, and further research is critical to fully elucidate the impact of HPAIV H5N1 in pregnant pinnipeds, as the current understanding is hindered by the limited data available. This is particularly significant in regions reporting high rates of abortions, where understanding the role of HPAI could help clarify its impact on reproductive health and population dynamics.

Our data confirmed viral infection in fetal organs including the lungs, heart, kidney, and thymus, while other organs couldńt be tested due to autolysis. Despite the extensive presence of infected cells in the lungs and heart, no associated lesions were observed, which is likely attributable to the immaturity of the fetal immune system. In contrast, in a full-term pup, widespread viral invasion of cardiac and pulmonary tissue would likely lead to severe cardiorespiratory failure with fatal outcomes. Moreover, the virus’s impact on the thymus induces atrophy and disrupts T lymphocyte development, resulting in severe lymphopenia and significantly impairs the immunocompetence of affected individuals at birth (52, 53, 54). Our findings are crucial for understanding the impacts of infection in pregnant females and their offspring, which may vary depending on the stage of gestation. Spontaneous abortions are likely to occur if transplacental infection takes place during early gestation when the fetus is not viable, as observed in cases of SASLs in Argentina. Conversely, perinatal mortality due to multiorgan failure may occur if the virus infects full-term pregnant females. This scenario could account for the high mortality rate observed among SES newborns at Peninsula Valdes during the 2023 breeding season (9, 12).

To our knowledge, this study represents the first pathological description of fatal lesions associated with HPAI H5N1, clade 2.3.4.4b in elephant seals. Fatal HPAI H5 infections have previously been documented in other seal species in the northern hemisphere, all characterized by neurological signs and CNS lesions (19, 21, 22). The severity and extent of the lesions observed in NEC 94 may provide a plausible explanation for the sudden and large-scale mortality reported among elephant seal pups at Peninsula Valdés. These findings remain speculative due to the limited sample size, which constrains our ability to draw definitive conclusions. The high mortality observed in pups may be influenced by their underdeveloped immune systems, increased susceptibility to HPAI infection, or a combination of these factors. In contrast, the relatively low number of adult SES carcasses recorded during the event suggests that this age group may have been less impacted by the virus. Alternatively, it is possible that some adults succumbed to infection after returning to sea, complicating efforts to comprehensively evaluate the effects of HPAI on older individuals. The scarcity of data highlights the urgent need for further research to determine whether SESs are more vulnerable to HPAI than other pinniped species. Understanding species-specific susceptibility is critical for assessing the virus’s overall impact on SES populations. Additional studies are needed to evaluate the broader consequences of HPAI on SES health, reproductive success, and long-term population viability, providing crucial information for conservation and management efforts.

Molecular analysis confirmed the virus as H5N1, clade 2.3.4.4b, with high sequence similarity to other HPAI HA segments reported across the Americas between 2022 and 2023 in various mammalian and avian species, including pinnipeds, as observed in this study. The analysis was conducted using FFPE tissue blocks, which represent a practical and versatile sample type. FFPE blocks can be stored at room temperature and easily shipped, making them an effective option for remote diagnostics and collaborative research. This method allows for the preservation of tissue morphology while enabling subsequent molecular analysis, thus facilitating a comprehensive understanding of the viral impact on pinniped health. In addition, molecular techniques applied to FFPE sections minimize the risk of human transmission of HPAI among researchers and laboratory workers, as the virus is effectively inactivated during fixation in formaldehyde.

This study was limited by a small sample size and the absence of systematic tissue sampling. Moreover, discrepancies were observed among the diagnostic tests performed on certain animals. For example, while NEC 85-R2 tested positive by PCR, with a Ct value of 34.4, indicative of a low viral load, it yielded negative results on IHC and exhibited minimal histopathological lesions. In contrast, NEC 83-R1 tested positive on IHC, showing a few scattered immunostained neurons and glial cells despite the presence of severe inflammatory lesions, yet was negative for the virus by PCR. This discrepancy could be explained by several factors, including the possibility that viral antigens were present in the tissue at the time of IHC analysis but were no longer detectable by PCR due to lower viral RNA presence in the sections used for PCR testing, which may have resulted from multiple cuts made between the two techniques. Consequently, these differences highlight the importance of employing a multistep diagnostic methodology to confirm cases of IAV. In addition, false negative results could arise from the non-uniform distribution of lesions and the partial sampling conducted on some animals, which limited the ability to conduct a thorough inspection and sampling of all organs. Further research employing larger sample sizes and systematic and standardized sampling protocols is essential for a more accurate diagnosis and understanding of HPAI in pinnipeds

Finally, HPAI H5N1 viruses represent a major public health threat due to their capacity to cross the species barrier and infect mammals. Recently, mammal-to-mammal transmission has been proposed based on mutation affecting genes related to mammal adaptation in strains isolated from South American pinnipeds (12, 31). These findings underscore the necessity for enhanced surveillance of HPAI in marine mammals and further investigation into the mechanisms of interspecies transmission and viral adaptation in non-avian hosts. Such research has broader implications for assessing potential risks to other mammalian species, including humans, particularly in environments with high levels of avian-mammalian interaction.

Our results are particularly relevant given the rapid evolution of HPAI viruses, highlighting the necessity for updated knowledge on the pathology associated with newer strains. Understanding the pathogenesis and effects of these recently circulating HPAI viruses in pinnipeds is essential for assessing their impact on these populations. This study provides a valuable perspective into key tissues affected by HPAI infection, offering a practical guide for targeted tissue collection that can enhance the accuracy and efficiency of HPAI diagnosis in pinnipeds. Our results demonstrate the value of pathology for understanding emerging diseases in wildlife, particularly relevant in the context of an emerging zoonotic pathogen.

## Acknowledgements

The authors thank the Chubut Coastal Wildlife Network for reporting pinniped strandings. We extend our gratitude to Lic. Víctor Fratto (REFAUNAR) and wildlife rangers María Cabrera, Marcia A. Schunk, Dulce M. Blanco, and Matías Tricase for their invaluable assistance during field sampling. We are especially grateful for their contributions in providing photographs and videos, which greatly enriched this study. Sample collection was conducted in full compliance with federal permits issued by the Dirección de Fauna y Flora Silvestre, Chubut, Argentina (Authorization N° 10/2023, DFyFSC).

## Author contributions

Study Design: C.F., A.F., E.S. Fieldwork and sample collection: C.F and P.A.A. Lab work and sample processing: C.F., A.C.R., M.A., E.S. Bioinformatics processing and data analysis: A.C.R., E.S. Manuscript preparation: C.F., A.F., M.A., D.L., E.S. Manuscript editing: all authors. Review and approval of final manuscript: all authors.

